# Comparative genomic analysis identifies great-ape-specific structural variants and their evolutionary significance

**DOI:** 10.1101/2023.03.16.533043

**Authors:** Bin Zhou, Yaoxi He, Yongjie Chen, Bing Su

## Abstract

During the origin of great apes about 14 million years ago, a series of phenotypic innovations emerged, such as the increased body size, the enlarged brain volume, the improved cognitive skill and the diversified diet. Yet, the genomic basis of these evolutionary changes remains unclear. Utilizing the high-quality genome assemblies of great apes (including human), gibbon and macaque, we conducted comparative genome analyses, and identified 15,885 great-ape-specific structural variants (GSSVs), including 8 coding GSSVs resulting in the creation of novel proteins (e.g. *ACAN* and *CMYA5*). Functional annotations of the GSSV-related genes revealed the enrichment of genes involved development and morphogenesis, especially neurogenesis and neural network formation, suggesting the potential role of GSSVs in shaping the great-ape-shared traits. Further dissection of the brain-related GSSVs show great-ape-specific changes of enhancer activities and gene expression in the brain, involving a group of GSSV-regulated genes (such as *NOL3*) that potentially contribute to the altered brain development and function in great apes. The presented data highlights the evolutionary role of structural variants in the phenotypic innovations during the origin of the great lineage.

## Introduction

The origin of the great ape lineage (the Hominidae family) about 14 million years ago represents an evolutionary leap in primates (Hill and Ward 1988; Pozzi et al. 2014), accompanied by a series of phenotypic innovations, including the increased body size (Smith and Jungers 1997), the enlarged brain volume (Barton and Venditti 2017; MacLeod et al. 2003), the improved cognitive abilities (Alba 2010) and the diversified diet (Chivers 1998). This evolutionary trend eventually leads to the origin of our own species. The living Hominidae family has four major lineages, including human (*Homo sapiens*), chimpanzee (*Pan troglodytes*), gorilla (*Gorilla gorilla*) and orangutan (*Pango pygmaeus*). It is of great interest to study the genetic basis of the great-ape-shared evolutionary traits that have made them a successful primate lineage, which in turn is informative to delineating the genetic mechanism of human origin.

Structural variants (SVs) are large genomic alterations (≥50 bp in length), such as deletions, insertions, copy number variations, inversions and translocations. They are widely distributed in the genomes of great ape species. For example, when comparing human and chimpanzee, the divergence at single nucleotide change is 1.23%, whereas the divergence of SVs reaches ∼ 3% of the genome (Suntsova and Buzdin 2020). Functionally, SVs are expected to have more impact than SNVs (single nucleotide variants). Previous studies have shown that in the human genome, SVs are predicted more harmful than SNVs (Abel et al. 2020), and more likely affect the expression of a gene than SNVs (Chiang et al. 2017). Specifically, SVs can affect the molecular function, cellular process, regulatory function, chromatin 3D structure, and transcriptional function of the organism (Spielmann, Lupiáñez, and Mundlos 2018; Weischenfeldt et al. 2013). As a major form of genetic variations, SVs also contribute to phenotypic diversity of organisms (Patel et al. 2014; Stankiewicz and Lupski 2010). However, previously, owing to the poor quality of the great ape genome assemblies (mostly based on next-generation sequencing, NGS), it has been difficult to systematically identify SVs and study their roles in phenotypic evolution.

Fortunately, the long-read sequencing and multiplatform scaffoldings are widely used in constructing high-quality reference genomes. For great apes, we now have high-quality genomes covering all major great ape lineages (Gordon et al. 2016; Kronenberg et al. 2018), providing a great opportunity to analyze SVs and their phenotypic relevance during primate evolution.

Here, we used the published high-quality genomes of human, chimpanzee, gorilla, orangutan, gibbon and macaque, and we identified 15,885 great-ape-specific SVs (GSSVs) through comparative genomic analyses. In particular, we report the potentially functional SVs that may contribute to the major phenotypic changes of great apes, especially the enlarged brains and the improved cognitive skills, which will lead to a better understanding of great ape evolution and human origin.

## Results

### Identification of the great-ape-specific SVs

The high-quality genomes of the great ape species (human, chimpanzee, gorilla and orangutan) were obtained from the published studies (Gordon et al. 2016; Kronenberg et al. 2018) (**Table S1**). To search for the great-ape-specific SVs (GSSVs), we also included the high-quality genomes of two outgroup primates: white-cheeked gibbon (*Nomascus leucogenys*) (NCBI accession number: GCF_006542625.1) and rhesus macaque (*Macaca mulatta*) (Warren et al. 2020) (**Figure 1A**).

**Figure 1.**
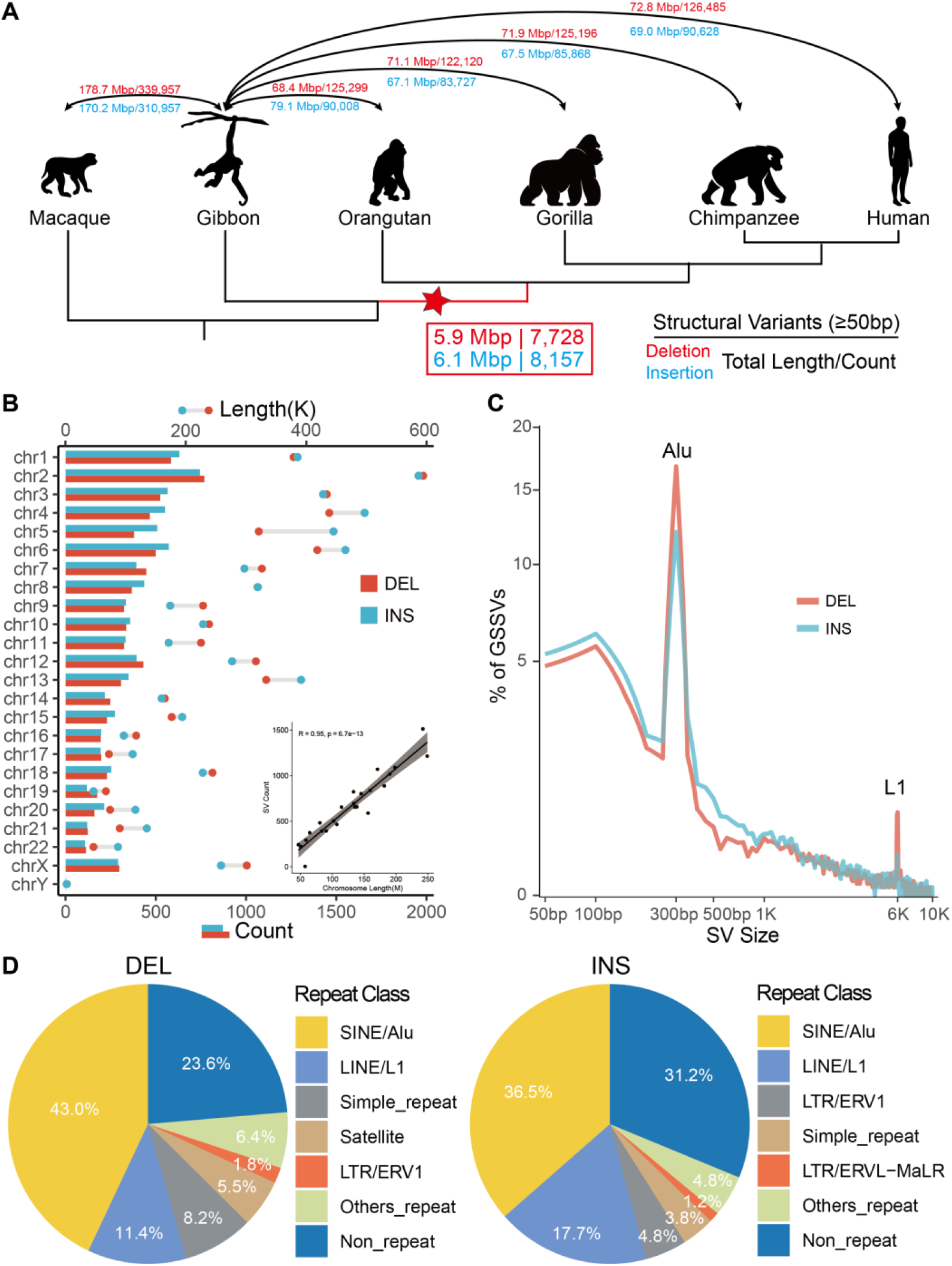
Identification and annotation of great-ape-specific SVs (GSSVs). (**A**) The cladogram indicates the phylogenetic relationship among the analyzed primate species. We used the high-quality genomes of four great ape species, one lesser ape species and one Old World monkey species. The numbers of SVs and their genomic lengths based on assembly comparisons are indicated. Deletions are labeled in red, and insertions in blue. The number in the red box indicates the identified 15,885 GSSVs, including 7,728 deletions and 8,157 insertions. (**B**) Chromosomal distribution of the identified GSSVs. The number of SVs is tightly correlated with the length of the chromosome, as indicated by the Pearson’s correlation (R2=0.95, P=6.7e-13). (**C**) The size distribution of GSSVs. The two peaks at 300 bp and 6 Kbp indicate the Alu and L1 elements, respectively. (**D**) Annotation of GSSVs located in the repeat regions of the genome, where SINEs (such as Alu) and LINEs (such as L1) compose of the great majority of the identified GSSVs.

Considering the methodological challenge for detecting SVs cross species with large genetic divergence (genome sequence identity < 95%), largely due to the difficulty of identifying syntenic blocks in the genomes (He et al. 2019), we assessed the two commonly used methods of detecting SVs based on an assembly-to-assembly strategy, including *smartie-sv* (Chaisson et al. 2015; Kronenberg et al. 2018) and *minimap2* (Feng and Li 2021; Li 2018). Based on the comparison of the callsets and manual curations, we found *minimap2* performed significantly better than *smartie-sv* in detecting SVs among distantly-related species (**Figure S1**). We therefore employed *minimap2* in the following analyses.

We first conducted pairwise genome comparisons between gibbon and the four great ape genome assemblies (human, chimpanzee, gorilla and orangutan) (Kronenberg et al. 2018). We then took the overlapped SVs among the four SV sets as the divergent SV set between gibbon and great apes (>50% reciprocal overlapping of SV length is required for an overlapped SV). Using the genome assembly of rhesus macaque (Warren et al. 2020), we further filtered out the SVs occurred in the gibbon lineage, which eventually gave rise to the set of GSSVs (see Methods for technical details). Totally, we detected 23,300 candidate GSSVs, including 14,451 deletions (DELs) and 8,849 insertions (INSs). Given the complex nature of SVs among deeply diverged species, to remove false positives, we further conducted manual curation of all the candidate GSSVs by local sequence alignments. Finally, a total of 15,885 GSSVs passed the curation, including 7,728 DELs and 8,157 INSs (**Figure 1A, Figure S2A**).

These GSSVs account for 12.0 Mbp genomic sequences (on average 0.44% of the great ape genomes), with fragment lengths ranging from 50 bp to 19 Kbp. They are randomly distributed in the genome, and the SV counts in each chromosome are positively correlated with the chromosome length (R = 0.95, *P* = 6.7e-13, Pearson correlation test; **Figure 1B, Figure S2B**), and 17.6% GSSVs (2,791) are longer than 1Kbp. As expected, we observed two peaks around 300bp and 6Kbp in the length distribution of the GSSVs, and the majority of them are Alu and L1 elements, respectively (**Figure 1C**). For repeat annotation, 76.4% DELs and 68.8% INSs are composed of repeat elements, such as SINEs, LINEs and LTRs (**Figure 1D**), suggesting that repeat elements are likely the key drivers for the generation of GSSVs.

### Functional implications of the great-ape-specific SVs

To search for GSSVs with potential functional consequences, we further classified the 15,885 GSSVs into two sets based on a range of criteria used in the aforementioned manual check (**Methods**), resulting in the set of 6,574 high-confident GSSVs (HC-GSSVs) and the set of 9,311 complex SVs (**Figure 2A**) (**Method**).

**Figure 2.**
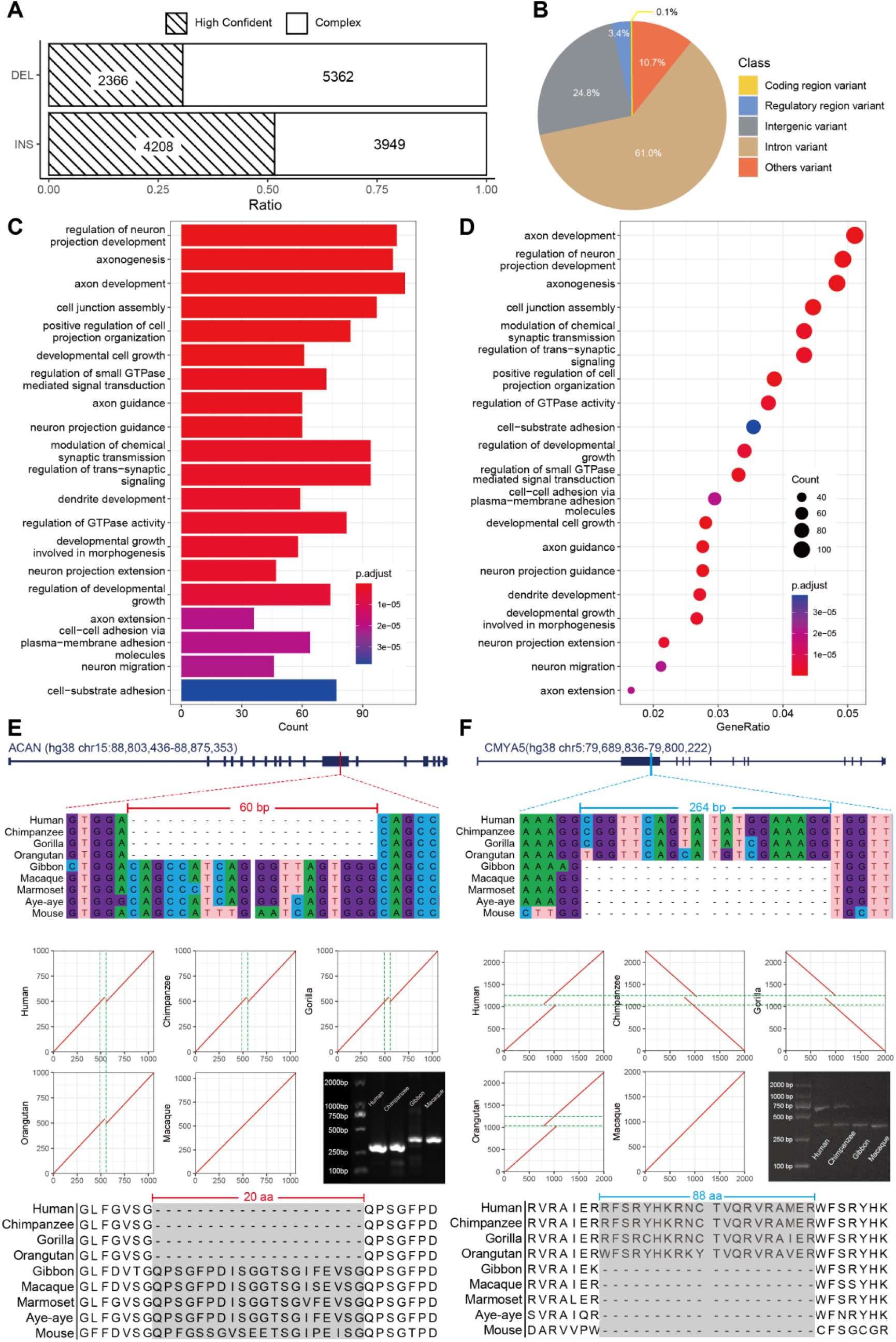
Functional annotations of the identified high-confident GSSVs (HC-GSSVs). (**A**) The ratios of the high-confident and the complex GSSVs after manual curation. (**B**) The pie plot shows the functional annotations of HC-GSSVs using the VEP tool. (**C-D**) The results of GO enrichment analysis of the 2,353 HC-GSSV-related genes. (**E**) A 60-bp deletion located in exon-12 of *ACAN*, a gene related to skeletal development. (**F**) A 264-bp insertion located in exon-2 of *CMYA5*, a well-known gene related to schizophrenia. The local sequence alignments (DNA and protein sequences) among mammalian species, together with the MUMmer plots showing the locations of the GSSVs are presented.

Our downstream analyses were focused on the HC-GSSV set containing 2,366 DELs and 4,208 INSs. We first conducted PCR validation of 10 DELs and 10 INSs randomly selected from the HC-GSSV set, and 19 of them were validated as true GSSVs, and only one failed because of nonspecific amplification of the target region (**Figure S3**). Accordingly, the HC-GSSV set can be considered a conservative set of GSSVs.

We annotated the identified HC-GSSVs using the VEP (Variant Effect Predictor) tool according to the human GRCh38 coordinates (**Methods**). The majority of HC-GSSVs (85.8%) are located in the intronic (61.0%) or the intergenic regions (24.8%), and 10.7% of them are located in the other regions (e.g. the 3’ or 5’ UTR regions). Notably, there are 3.4% (222 GSSVs) located in the known regulatory regions such as enhancers and promotors, an implication of their potential roles in gene expression regulation.

Next, we conducted functional enrichment analysis for the 2,353 HC-GSSV-related genes (those genes overlapped with the HC-GSSVs, see **Methods** for details) (**Table S8**). The enriched functional terms are related to the known traits of the great ape lineage, such as developmental growth involved in morphogenesis (body size) and axonogenesis (brain). Remarkably, more than half of the enriched terms are related to brain development and function. The top terms include axon development, neurogenesis, neural projection, and neuronal differentiation (**Figure 2C and D**). These results indicate a potential role of HC-GSSVs in shaping the great-ape-specific phenotypic traits, especially the central nervous system.

In particular, we found 8 coding HC-GSSVs (2 DELs and 6 INSs) (**Table 1 and Figure 2B, Figure S4**). Although several of them are located in exons and were predicted causing protein sequence changes, none of them actually result in truncated proteins (**Table 1**). Among these GSSV-related genes, *POU5F1B* is a great-ape-specific novel gene caused by the retro-position of its mother copy (*OCT4*) on chromosome 8 (a 1,412-bp INS specifically occurred in the great-ape lineage), confirming the previous report (Simó-Riudalbas et al. 2022). Initially, it was thought to be a pseudogene of *OCT4* with 95% sequence similarity. However, there have been reports suggesting its susceptibility in tumors (Rafnar et al. 2014; Zanke et al. 2007), and it likely acts as a transcriptional activator to promote cancer cell proliferation (Pan et al. 2018, 1; Panagopoulos et al. 2008). In addition, *POU5F1B* is implicated in the regulation of *OCT4* activity (Suo et al. 2005). The functional role of *POU5F1B* in the evolution of great apes is yet to be understood.

**Table 1.** The 8 HC-GSSVs located in the gene-coding regions.

| Position<sup>#</sup> | Length (bp) | Type | Gene | Consequence<sup>*</sup> | Gene function or disease |
| --- | --- | --- | --- | --- | --- |
| Chr2:232,330,767-232,330,767 | 186 | DEL | *DIS3L2* | c,d | RNA degradation |
| Chr15:88,857,790-88,857,790 | 60 | DEL | *ACAN* | d | skeletal development |
| Chr5:79,736,810-79,736,810 | 264 | INS | *CMYA5* | e | schizophrenia |
| Chr15:81,135,350-81,135,983 | 633 | INS | *CFAP161* | a,f | ciliary motion |
| Chr11:71,961,971-71,962,270 | 299 | INS | *RNF121* | a,g,h | regulation of cell cycle, signal transduction |
| Chr8:127,415,807-127,417,219 | 1412 | INS | *POU5F1B* | b,h | weak transcriptional activator |
| Chr10:77,994,516-77,999,310 | 4794 | INS | *POLR3A* | a,f,g,h | spastic ataxia |
| ChrX:14,915,424-14,915,588 | 164 | INS | *MOSPD2* | a,g,h | intracellular exchanges and communication |

Note: <sup>#</sup>the coordinates of the human GRCh38; <sup>\*</sup>a) coding\_sequence; b): start\_lost; c) stop\_gained; d) inframe\_insertion; e) inframe\_deletion; f) splice\_donor; g) splice\_acceptor; h) UTR

Interestingly, one coding HC-GSSV, a 60-bp DEL, is located in exon-12 of *ACAN*. This gene is involved in skeletal development (**Figure 2E**). It is one of the major components of cartilage, and binds to hyaluronan and links protein to form huge aggregates, a hydrated gel-like structure with resistibility to compression and deformation in joints (Watanabe, Yamada, and Kimata 1998). Many studies have shown that mutations in *ACAN* can lead to short stature and poor bone development (Hu et al. 2017; Wei et al. 2021). Besides, as the main component of the neural extracellular matrix, *ACAN* is also expressed in the brain (Morawski et al. 2012). We speculate that the deletion of 20 amino-acids (due to the 60-bp DEL) in the binding domain of the free sugar chains of *ACAN* may affect bone development in great apes, which might be related the increased body size during the origin of the great ape lineage.

The other potentially important case is a 264-bp INS located in exon-2 of *CMYA5*, resulting in an 88 amino-acid insertion (**Figure 2F**). Previous GWAS studies have found that variants in *CMYA5* are associated with schizophrenia (Chen et al. 2011; Hoya et al. 2018), implying that this HC-GSSV in *CMYA5* may play a role in the brain evolution of great ape species. Another interesting case is the 186-bp DEL in *DIS3L2*. This gene is mainly related to RNA degradation, and mutations in this gene can cause Perlman syndrome, characterized by a larger head size and developmental delay (Morris, Astuti, and Maher 2013), providing a hint of this GSSV in the great-ape-specific pattern of brain/head development.

The remaining 4 HC-GSSVs are not associated with the known great-ape-specific traits based on current knowledge. *CFAP161* is associated with cilia movement (Austin-Tse et al. 2013), while *RNF121* is involved in the regulation of cell cycle, signal transduction, genomic integrity under hypoxia, and metastasis of cancer cells (Gao et al. 2017; Lee et al. 2018). *POLR3A* is associated with spastic ataxia, and the best recognized phenotype is cerebellar ataxia (Infante et al. 2020), and *MOSPD2* is involved in intracellular exchange and communication (Di Mattia et al. 2018). Whether and how these HC-GSSVs contribute to the origin and evolution of great apes are yet to be explored (**Figure S4**).

### The regulatory HC-GSSVs with potential functions in the brain

Compared to gibbons and Old World monkeys, the most crucial changes that occurred during the origin of the great apes include the increase in body size, the expansion of brain volume, and the improvement of cognitive ability. As aforementioned, the GO enrichment analysis shows that most of the GSSVs-associated genes are involved in the central nervous system, and the majority of the HC-GSSVs were mapped to the intronic or intergenic regions, suggesting that their functional impact may relate to gene expression regulation. It has been postulated a few decades ago that the differences between human and chimpanzee are mostly caused by gene regulation changes rather than by alterations in their protein-coding sequences (King and Wilson 1975), and a recent study also reached the same conclusion (Suntsova and Buzdin 2020).

With the use of the published brain ChIP-seq (H3K27ac) and RNA-seq data from human, chimpanzee, and rhesus macaque (Sousa et al. 2017; Vermunt et al. 2016), we identified 105 HC-GSSVs that overlapped with105 human-chimpanzee specific cis-regulatory elements (CREs). Among the 105 CREs, there are 186 associated genes (genes within the 500 Kbp downstream and upstream of CRE) showing human-chimpanzee specific expression changes compared with rhesus macaque, and the change direction of gene expression is consistent with the H3K27ac signals (indication of CRE activity). These 105 CREs include 36 DEL-containing CREs associated with 62 nearby genes, and 69 INS-containing CREs associated with 126 genes (two genes are present in both gene sets) (**Table S14**).

We firstly ranked the 43 up-regulated genes (human-chimp vs. macaque) by log2 fold changes of gene expression. The top 5 up-regulated genes include *GGTA1, ZGLP1, C20orf202, NOL3*, and *MAP1LC3B* (**Figure 3A**). Of these genes, *NOL3* is closely associated with neurodevelopment. Markedly, *NOL3* also ranks the top gene in the occipital pole (OP) of the cortex (**Figure 3B**). Previous studies have shown that *NOL3* is related to abnormal neural potential, and the absence of *NOL3* causes excessive excitation (Russell et al. 2012). *NOL3* is 42.9 Kbp away from the nearest human-chimpanzee specific CRE, and a 297-bp INS is located in this CRE. The H3K27ac peak map shows that human and chimpanzee have higher peaks than that of macaque (P-value < 0.05, Welch’s two-tailed unpaired t test). The multiple sequence alignment shows that the genomic region containing this INS is conserved in primates (**Figure S5A**), and the 297-bp INS likely causes a CRE activity change, creating a human-chimpanzee specific enhancer that regulates *NOL3* expression.

**Figure 3.**
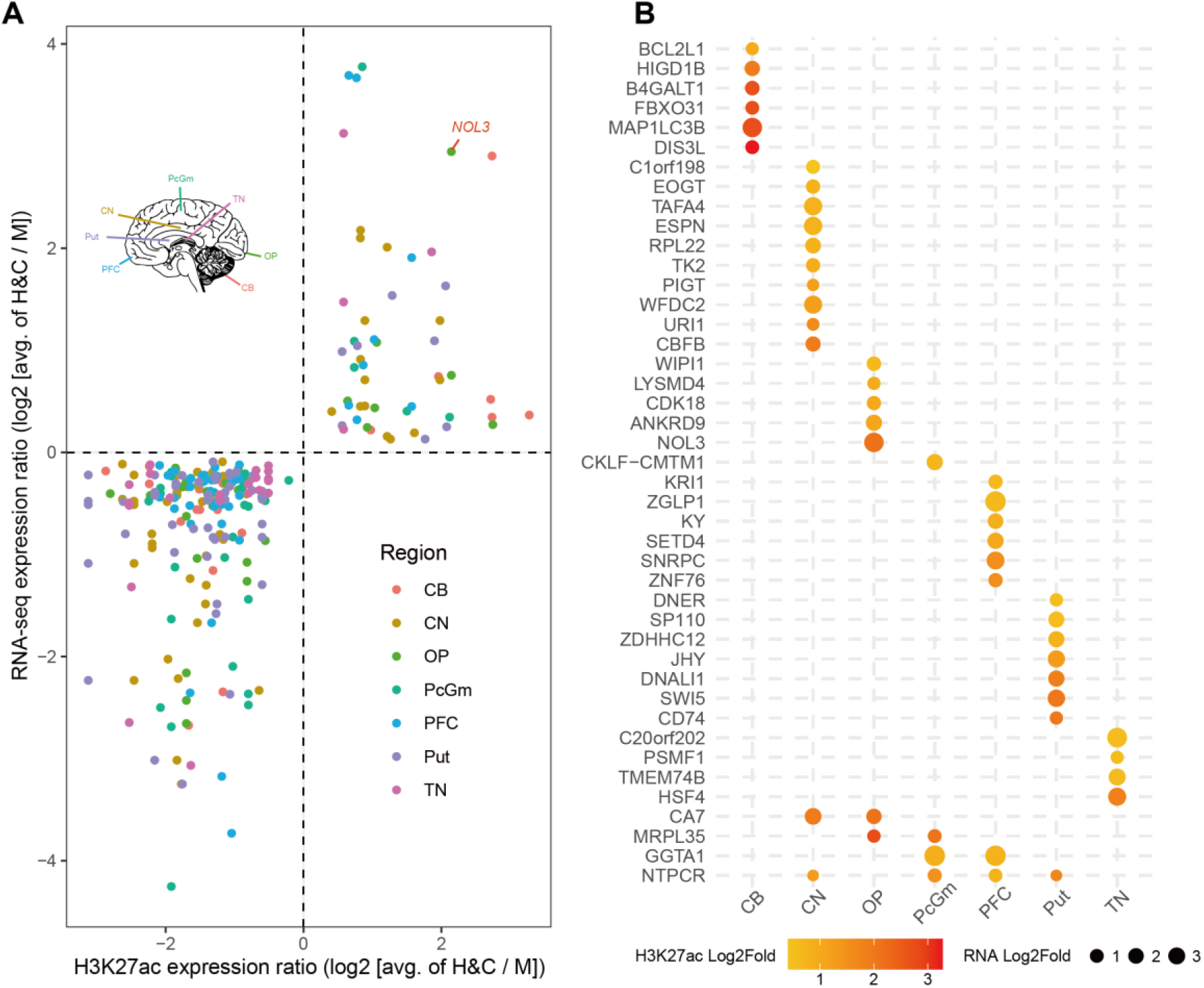
The HC-GSSVs located in the cis-regulatory elements (CREs) of the brain. (**A**) The HC-GSSV-related genes showing human-chimp specific CRE activity (by the H3K27ac signals) and gene expression changes (by RNA-seq) compared to macaque across 7 brain regions, including cerebellum (CB), caudate nucleus (CN), occipital pole (OP), precentral gyrus (PcGm), prefrontal cortex (PFC), putamen (Put) and thalamic nuclei (TN). (**B**) Ranking of the 43 up-regulated genes based on gene expression and the H3K27ac signals.

In addition, using the FIMO tool (Grant, Bailey, and Noble 2011), we performed transcription factor (TF) enrichment analysis of the homologous CRE regions, and we identified 78 human-chimpanzee-specific TFs (**Figure S5B**) with 27 of them located in the HC-GSSV regions. The strongest signal (by the FIMO score) is *ZNF460* (**Figure 4C, Table S15**), a TF mainly expressed in the brain (**Figure S5D**), though its function is largely unknown.

**Figure 4.**
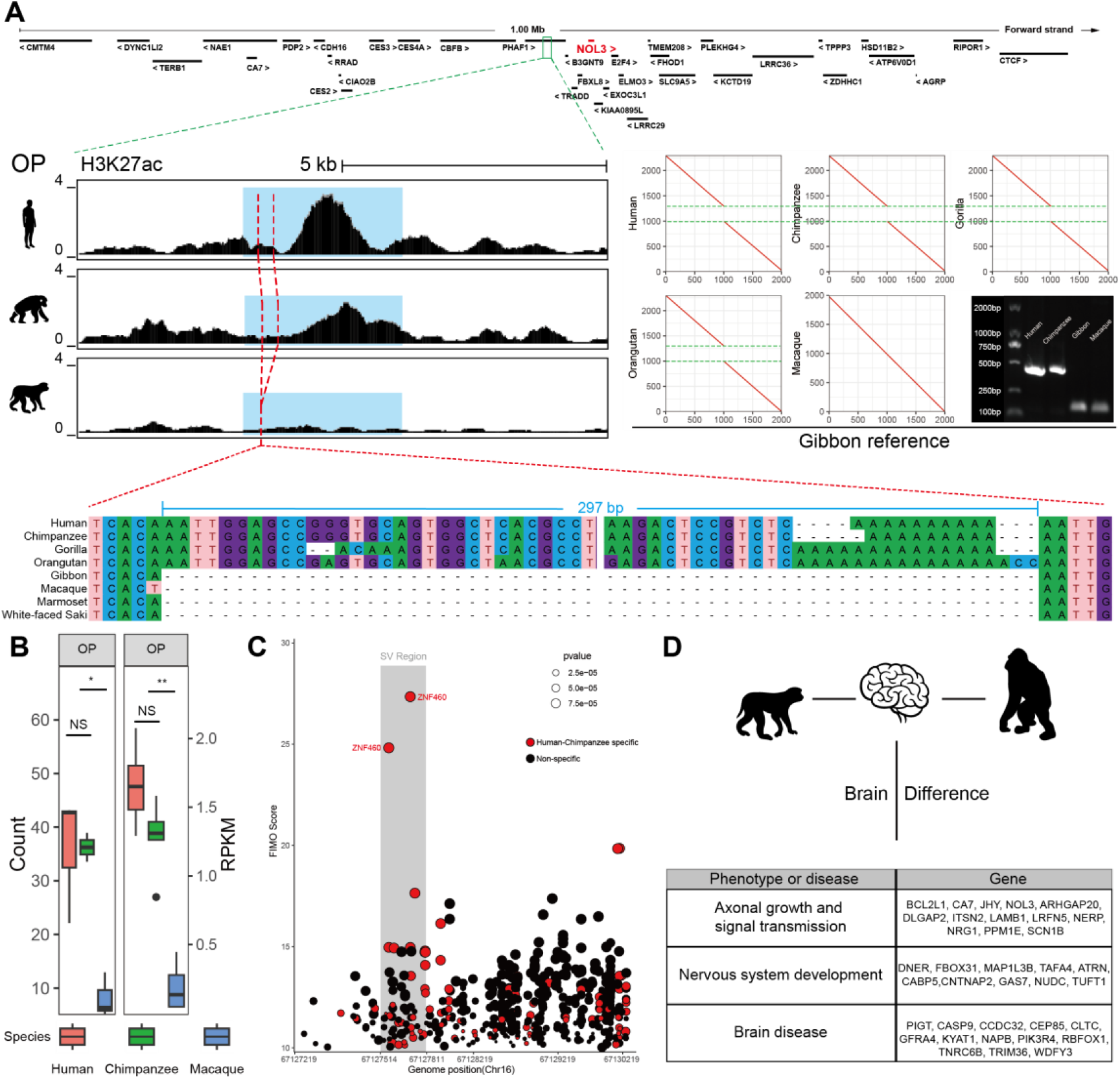
Overview of GSSV occurred in *NOL3*. (**A**) A 297-bp INS located in a Human-Chimpanzee specific CRE (blue region) which the nearest Human-Chimpanzee specific gene is *NOL3*. The left plot shows the difference of H3K27ac signal in occipital pole (OP) region among human, chimpanzee and rhesus monkey; the right plot shows the MUMmer plot of the 297-bp INS between great apes and gibbon; and bottom plot shows multiple sequence alignments of representative primate species. (**B**) The comparisons of normalized read count of H3K27ac signal of CRE (left) and expression (right) of *NOL3*. NS, not significant; *, *P*<0.05, **, *P*<0.01. (**C**) TF enrichment of CREs. The CREs with Human-Chimpanzee specific TF binding sites are denoted in red scatters, and two *ZNF460* binding sites are labeled which have the highest FIMO score and located in SV region (grey region). (**D**) Summary of brain-related genes with human-chimp specific CREs and overlapped to identified GSSVs.

For the 143 down-regulated genes (**Figure S6**), there are also worth-noting cases, such as *LRFN5*, 21.4 Kbp away from the nearest human-chimpanzee-specific CRE, and a 508-bp INS is located in this CRE (**Figure S7**). As a synaptic adhesion molecule implicated in autism, *LRFN5* can induce presynaptic differentiation through binding to the LAR family receptor protein tyrosine phosphatases (LAR-RPTPs) that have been highlighted as presynaptic hubs for synapse formation (Goto-Ito et al. 2018; Lin et al. 2018).

The brains of great apes are structurally and functionally more complex than those of gibbons and macaques. We summarized the annotated functions of all the identified HC-GSSV-related genes showing human-chimpanzee-specific expression changes in the brain. They are mainly involved in three functional aspects, including synapse formation and signal transmission, nervous system development (such as neural migration and neural differentiation), and various brain diseases caused by gene mutations (intellectual disability, microcephaly, schizophrenia, Parkinson, etc.) (**Figure 4D**).

Taken together, we identified a set of genes related to HC-GSSVs which are associated with great ape brain function. These HC-GSSVs could serve as a resource to study the genetic basis of brain structure/function change that emerged during the origin of the great ape lineage. However, owing to the lack of ENCODE data of the correspondent tissues in most of the non-human primate species, the speculated regulatory roles of these HC-GSSVs are yet to be validated.

## Discussion

Based on the high-quality great ape assemblies, we identified 15,885 great-ape-specific SV by cross-species comparison. Through further manual curation with a stringent filtering, we report 6,574 high-confidence GSSVs that overlap with 2,353 genes. These HC-GSSV-related genes show functional connections with the great-ape-specific traits, such as body size and brain. Markedly, many of the enriched functional terms of these genes are related to brain development and function, implying an important role of structural variants in shaping the central nervous system of great apes during evolution. In particular, we report 8 coding GSSVs that lead to the generation of novel proteins during the origin of the great ape lineage, and some of these great-ape-specific proteins (such as *ACAN* and *CMYA5*) are involved in bone development and brain function, providing clues to the genetic basis of the great-ape-shared phenotypic innovations.

Previous studies had suggested that the body size or weight of great apes are significantly larger than lesser apes and Old World monkey (Smith and Jungers 1997; Wheatley 1987; Zihlman and Bolter 2015; Zihlman and McFarland 2000). In the HC-GSSVs set, we found a 60-bp DEL leading to a 20 amino-acids deletion of *ACAN*, a gene related to bone development. Previous genetic studies in humans reported that the *ACAN* mutations could cause short stature and spinal disease (Hu et al. 2017; Wei et al. 2021). Hence, this SV-induced novel *ACAN* protein may play a potential role in body size evolution of great apes. Another important phenotypic innovation in the great ape lineage involves the brain. Accordingly, we identified many HC-GSSVs associated with brain development and function. For example, a 264-bp INS changes the coding sequence of *CMYA5*, and previous human studies suggest its involvement in schizophrenia (Chen et al. 2011; Hoya et al. 2018).

Besides of the 8 coding GSSVs, the great majority of the identified GSSVs are located in the non-coding regions of the genome, which presumably contribute to gene expression regulation. One notable example is a 297-bp INS that influences the enhancer of *NOL3*, a gene related with abnormal neural potential.

During primate evolution, there have been two leaps of brain volume enlargement, one in the common ancestor of the great ape lineage and the other in the human lineage (Holloway et al. 2009). Consequently, the brain volume of great apes is much larger than lesser apes. The larger brain means an improved cognition, because of the higher relative cortex volume and neuron packing density (NPD). Likewise, information processing capacity becomes higher due to short inter-neuronal distance and high axonal conduction velocity (Roth and Dicke 2012). Thanks to the published brain epigenomic data that include human, chimpanzee and macaque (Sousa et al. 2017; Vermunt et al. 2016), we were able to search for the GSSVs located in the CREs and to infer their potential influences on gene expression regulation. We found many genes affected by GSSVs are related with brain size, such as *WDFY3, CCDC32* and *CLTC*, all of which are involved in microcephaly. More importantly, these genes are also associated with intellectual disability and developmental delay (Abdalla et al. 2022; DeMari et al. 2016; Le Duc et al. 2019; Nabais Sá et al. 2020). Given the more complex cortex structure of great apes, we found several GSSV-related genes (*FBXO31* and *NUDC*) acting on neuronal differentiation and migration, which may help form the complex neural network in the brain. Consistently, several GSSV-related genes (*BCL2L1, CA7* and *LRFN5*) are involved in axonal growth and signal transmission, highlighting their potential roles in the higher information processing capacity of great apes. Together, the enriched GSSVs and the target genes for brain development and function indicate the evolutionary significance of structural variants that may contribute to the larger brain and higher cognitive abilities of great apes. Future functional experiments are warranted to reveal the molecular and developmental mechanisms underlying the bigger and smarter brains of great apes.

In conclusion, given the challenge of studying SVs among distantly-related species, in this study, we identified the great-ape-specific SVs, providing a useful resource for understanding the genetic basis of phenotypic innovations in the evolution of great apes.

## Method

### Comparing *Minimap2* and *Smartie-sv*

We mapped each long-read assembled genome of the four great ape species to the gibbon assembly (NCBI accession number: GCF_006542625.1) and calling SVs by *Minimap2+paftools* and *Blasr+printgaps (Smartie-sv)* respectively. For the reverse calling, we called the SVs by mapping the gibbon genome back to each of the great ape genomes. We calculated the overlap between two methods and used the commmand “ shuf –n 100 SV_list.bed” and repeated 10 times to randomly select condidate SVs and futher validate the ture SVs rate by manual check (**Figure S1**).

### Identification of GSSV

Genome comparisons were performed using minimap2 (Li 2018). We mapped each long-read assembled genome of the four great ape species to the gibbon assembly (NCBI accession number: GCF_006542625.1), including human-GRCh38.p13 (V38), Chimpanzee-Clint_PTRv2 (CCP), Gorilla-Kamilah_v0 (GGO), Orangutan-Susie_PABv2 (PAB) and rhesus-Mmul_10 (RM10). Using gibbon genome as the reference genome, we mapped the genomes of great apes to the gibbon genome for SV calling, referred as the forward calling. For the reverse calling, we called the SVs by mapping the gibbon genome back to each of the great ape genomes. Then, we filtered the SVs by intersecting the two SV sets and obtained the SVs for each pair (**Table S2**). GSSVs were identified by taking the intersection of all the gibbon–great ape SV sets. Furthermore, to exclude the gibbon lineage-specific SVs, we used the published rhesus macaque genome to repeat the forward and reverse calling, and the intersected GSSVs set was taken as the final set of GSSVs (**Table S3, Table S4**).

### Repeat analysis

Transposable elements (TEs) were identified by using RepeatMasker (v4.0.9) to search against the known Repbase TE library (Repbase21.08). We identified the TE ratios of INS in the human genome and DEL in the gibbon genome, respectively (**Table S7**).

### GSSVs manual check and PCR validation

The sequences of the GSSV regions were extracted by the “bedtools getfasta” command, and sequence comparison was performed and plotted by MUMmer (NUCmer-3.1 and MUMmerplot-3.5). The candidate GSSVs with 1 Kbp upstream/downstream sequences were aligned to the gibbon genome using NUCmer. The GSSVs were classified into three categories based on manual check: (i) If the GSSV region and its 1 Kbp flanking sequences of macaque could be completely aligned to the corresponding gibbon GSSV region using NUCmer and MEGA-7.0.26 (MUSCLE under the default parameters), they were considered as orthologous between gibbon and macaque. These GSSVs were defined as “high-confident” GSSVs (HC-GSSV); (ii) If the GSSV region with 1 Kbp flanking sequences of macaque could only be partially aligned to the corresponding gibbon GSSV sequence, we classified these GSSVs as “complex” GSSVs; and (iii) If the potential GSSV region with 1 Kbp flanking sequence of macaque could be completely aligned to the corresponding gibbon GSSV sequences except for the SV regions, we inferred that these SVs are present in macaque, and we defined these GSSVs as “false” GSSVs. The detailed list is shown in **Figure S9**. For PCR and Sanger sequencing validation, we included one rhesus macaque, one white-cheeked gibbon, one chimpanzee and one human. The primers were designed by Primer Premier5 (**Table S17**). The PCR products were visualized by agarose gel electrophoresis to verify the lengths of GSSVs and for Sanger sequencing. All the presented SVs in this study were validated by both PCR and Sanger sequencing.

### GO enrichment analysis

By command “bedtools intersect –a SV_coord.bed –b Gene_coord.bed –wa –wb” to get the genes which intersect with SVs (**Table S8**) and futher do GO enrichment analysis by ClusterProfiler-4.6.0 (Wu et al. 2021).

### Annotation analysis for HC-GSSVs

HC-GSSVs annotation was performed using VEP with the GRCh38 coordinates (**Table S10, Table S11**).

### Identification of HC-GSSVs located in the great-ape-specific CREs

The published ChIP-seq data were used to identify the great-ape-specific CREs in seven brain regions that have correspondent H3K27ac and RNA-seq data (Sousa et al. 2017; Vermunt et al. 2016). We downloaded the 60,703 genomic regions with H3K27ac signals, which were generated from three humans, two chimpanzees and three rhesus monkeys in each of the seven brain regions, including prefrontal Cortex (PFC), precentral gyrus (PcGm), occipital pole (OP), caudate nucleus (CN), putamen (Put), cerebellum (CB), and thalamic nuclei (TN). We performed pair-wise comparison of the enhancer signals (the H3K27ac peaks) among the three species (macaque–human, macaque–chimpanzee, and human–chimpanzee), and we used the Welch two-tailed unpaired t test for statistical assessment. We defined an enhancer as an great-ape-specific CRE if this enhancer showed significant difference (P < 0.05) of the same direction in both the macaque–human and the macaque–chimpanzee comparisons while showing no difference (P > 0.05) in the human–chimpanzee comparison. According to this criterion, we obtained 18,105 human-chimpanzee specific CREs (**Table S12**), among which 105 of them are overlapped with the HC-GSSVs, involving 186 genes. To quantify the activities of these great-ape-specific CREs, we calculated the log2(fold-change) between great apes (the average of human and chimpanzee) and rhesus macaque (**Table S14**).

### Comparative gene expression analysis of seven brain regions

The gene expression data of adult brains of human, chimpanzee and macaque were obtained from the published data (Sousa et al. 2017), which contains samples from six humans, five chimpanzees and five macaques of 16 brain regions. To match the expression data with the ChIP-seq data, we used the data of 9 brain regions, including primary motor cortex (M1C), mediodorsal nucleus of the thalamus (MD), primary visual cortex (V1C), cerebellar cortex (CBC), striatum (STR), ventrolateral prefrontal cortex (VFC), orbital prefrontal cortex (OFC), dorsolateral prefrontal cortex (DFC), medial prefrontal cortex (MFC), and the matching relationships between the ChIP-seq and the gene expression data is shown in **Figure S7**. The Welch two-tailed paired t test was used in accessing expression differences between great apes and macaques and obtained 1,714 human-chimpanzee specific genes (**Table S13**).

### Transcription factor (TF) enrichment analysis

We used liftover to obtain the homologous sequences of the great-ape-specific CREs (chr16_67128753_hg38) in human, chimpanzee and macaque. Then, using FIMO, we predicted the enriched TFs on the sequence of each species, and screened out the human-chimpanzee specific TFs (**Table S15**).

## Data availability

All genome, epigenome and transcriptome data could be download in NCBI database and the code used in this study had submitted to github.

## Author contributions

B.S. designed the project; B.Z and Y.H. performed bioinformatics analyses, genotyping and sequencing experiments; B.S., Y.H. and B.Z. wrote the manuscript. All authors have discussed the results and read the manuscript.

## Competing interests

The authors declare no competing financial interests.

## Acknowledgements

This work has been supported by grants from the National Natural Science Foundation of China (U2002207 to B.S.), and the National Key Research and Development Program of China (2021ZD0200100 to B.S.).

## Supplementary Figures 1-9

**Figure S1.**
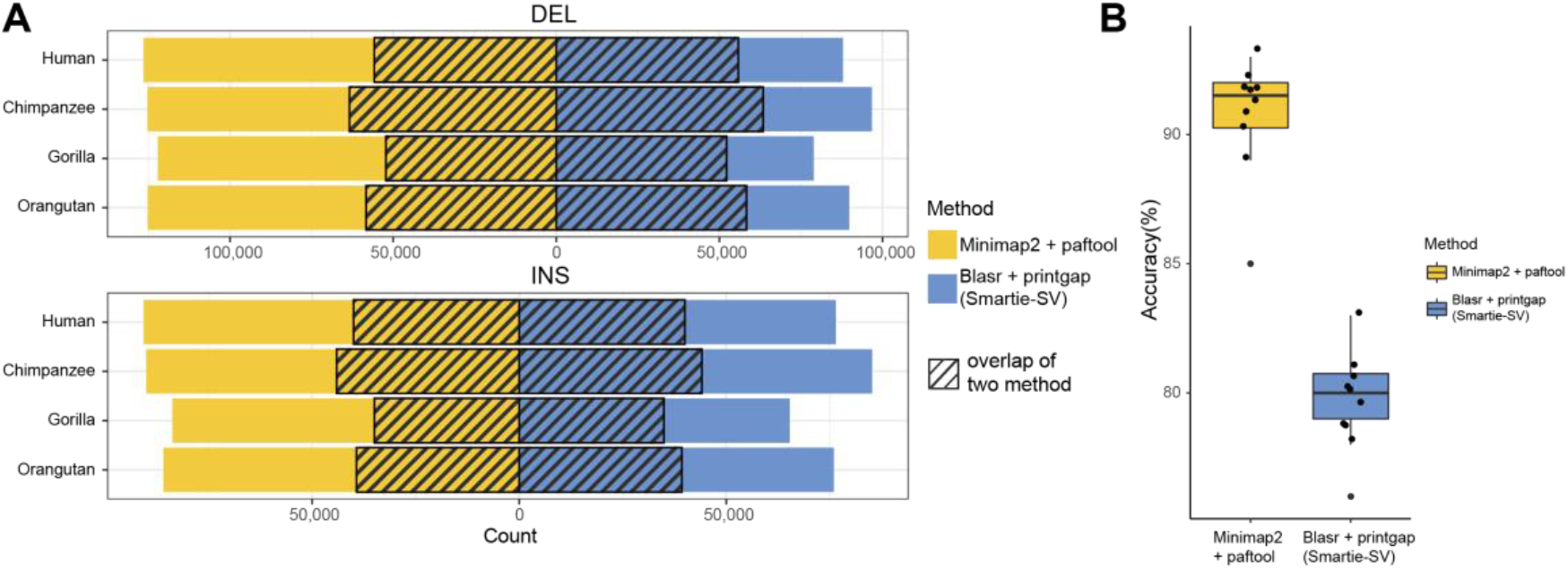
Comparison of the two SV mapping/calling methods. (A) The identified SVs by mapping four great ape genomes to the gibbon genome. The diagonal boxes indicate the overlapped SVs between the two methods. (B) Comparison of the accuracy between the two methods. We randomly selected 100 SVs in the SV set, and conducted manual check to calculate the rate of accuracy. The manual check was repeated 10 times.

**Figure S2.**
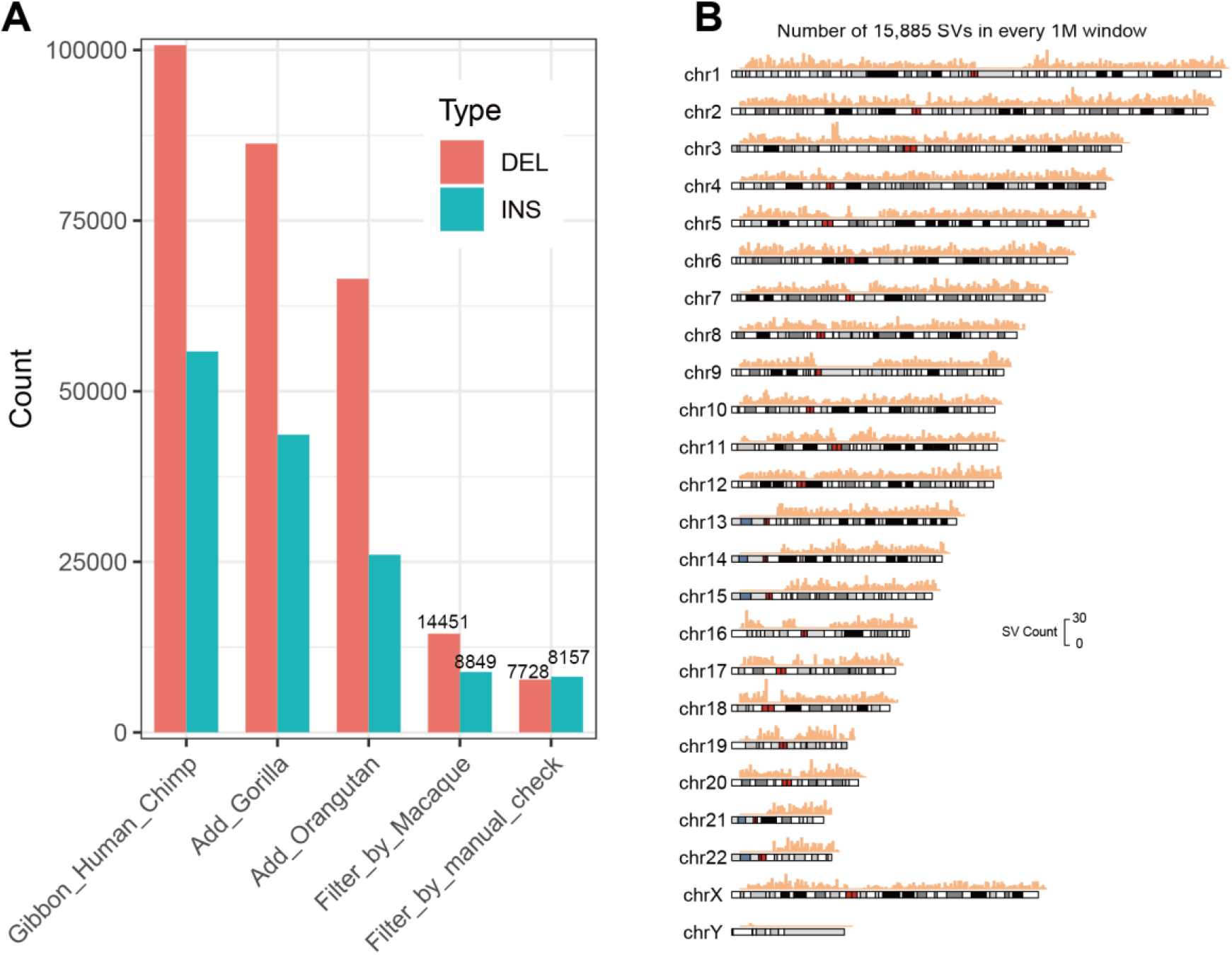
The filtering process of the SV numbers and the chromosomal distribution of the identified 15,885 GSSVs. (A) Th SV counts during the filtering processing that leads to the final set of 15,885 GSSVs, including 7,728 DELs and 8,157 INSs. (B) Chromosomal distribution of the 15,885 GSSVs in the 1Mbp windows (hg38).

**Figure S3.**
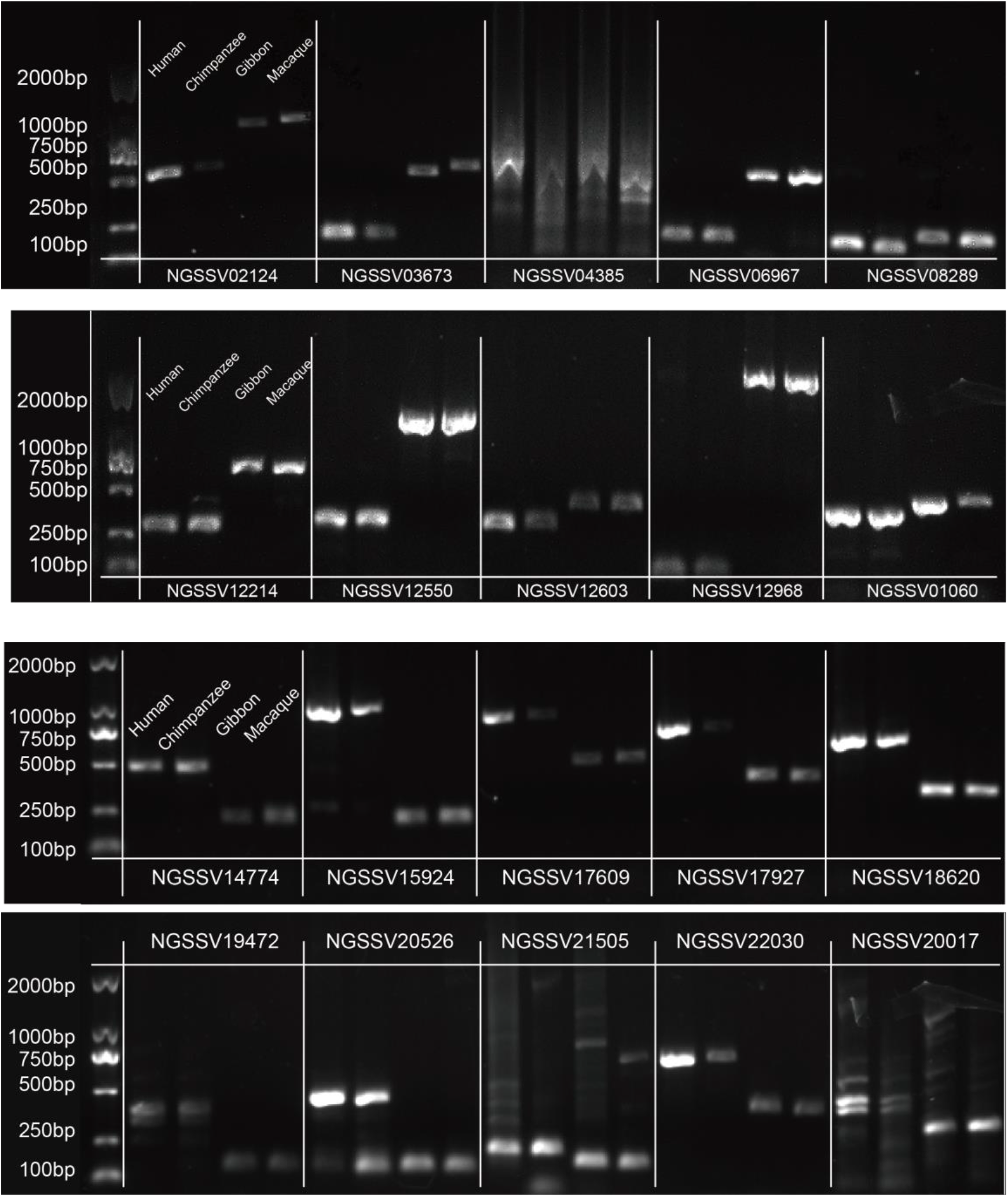
PCR validation of 10 DELs (top) and INSs (bottom). A total of 19 SVs were validated except for one DEL showing non-specific amplification.

**Figure S4.**
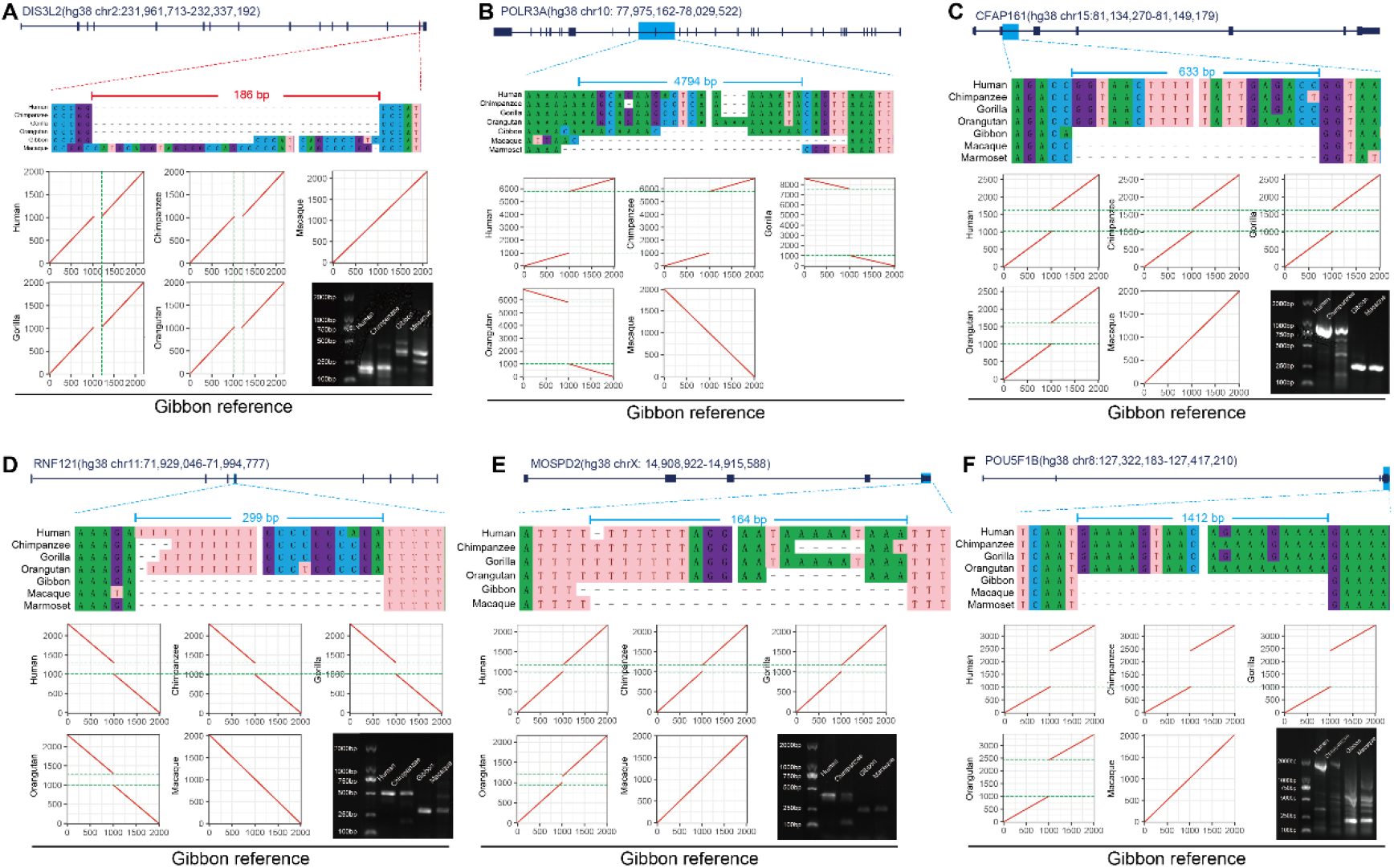
The data of 6 coding SVs. For each case, the SV position, the DNA sequence alignment and the MUMMER plot are presented. The POLR3A SV (4794bp insertion) was not PCR validated to the large SV size and its genomic region containing a large number of copy number variants (CNVs).

**Figure S5.**
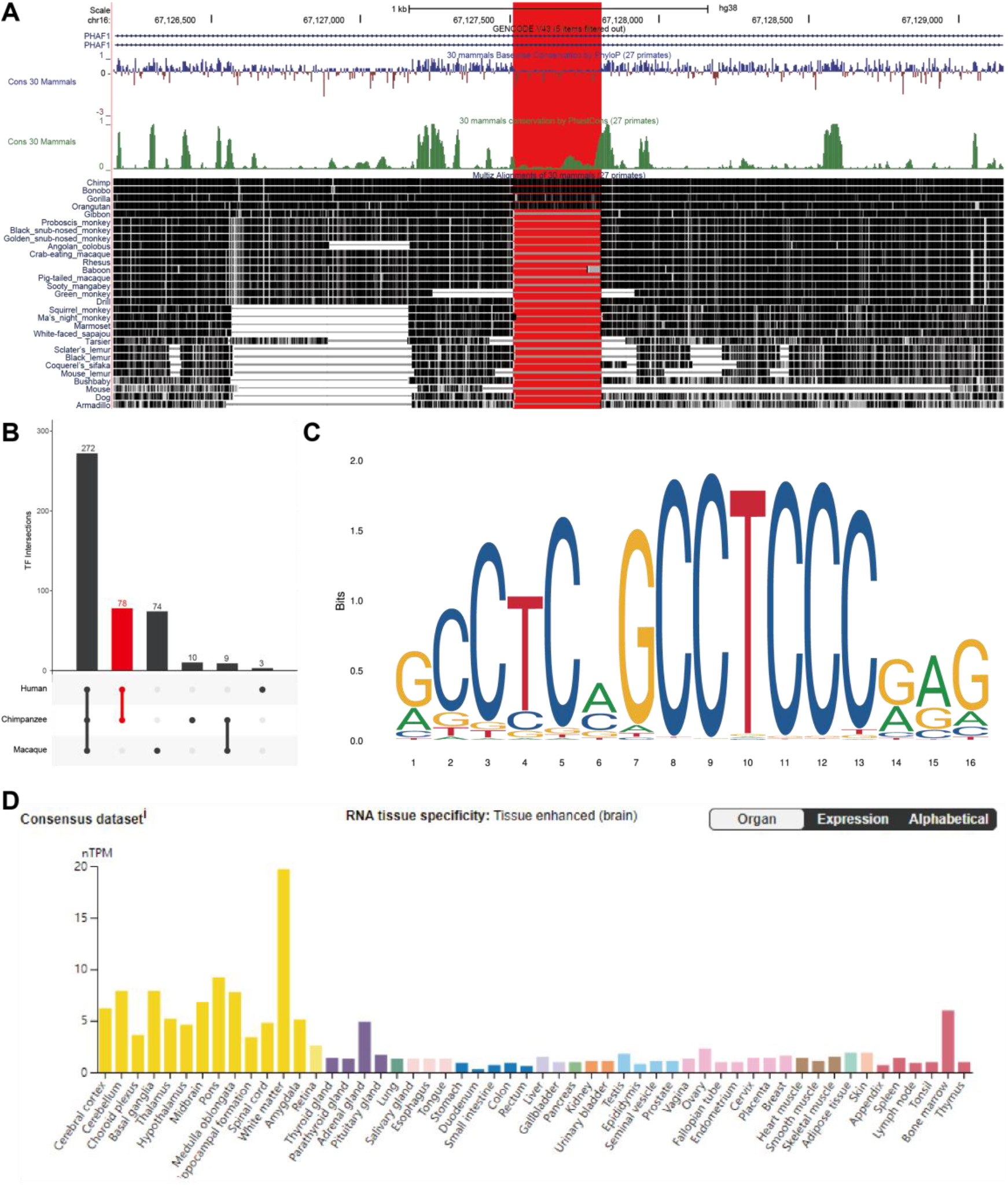
Sequence conservation of the HC-GSSV located genome region and TF enrichment of the SV-containing CRE. (A) The UCSC plot shows the sequence conservation of the genomic region containing the SV among 30 mammalian species (the red box highlights the SV region). (B) The 78 human-chimpanzee-specific TFs (red bar). (C) The binding sequence motif of the transcription factor ZNF460. (D) The mRNA expression levels of ZNF460 in all human tissues, and the brain shows the highest expression (yellow bar). The data is from The Human Protein Atlas (https://www.proteinatlas.org).

**Figure S6.**
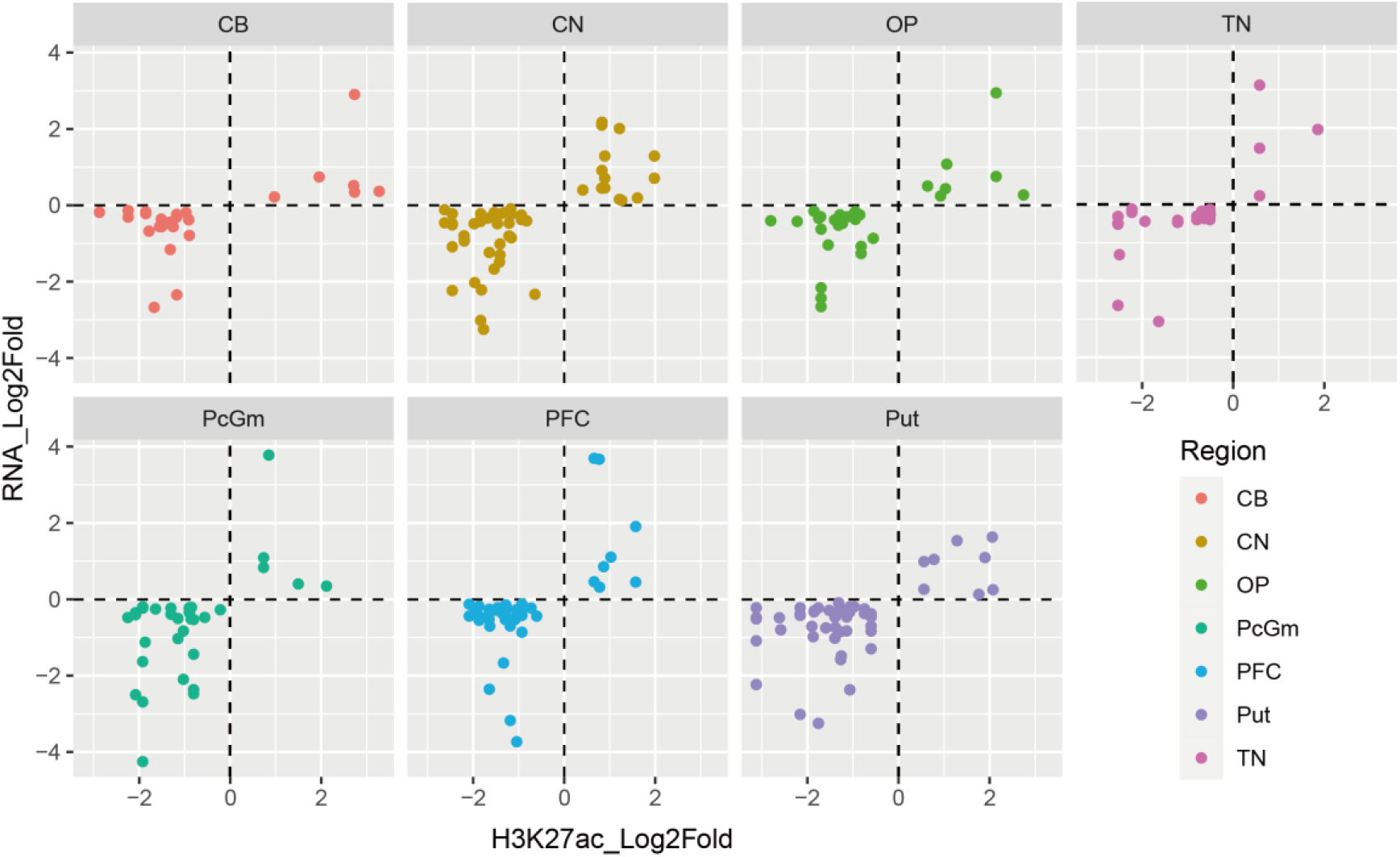
The up-regulated and the down-regulated genes in seven brain regions.

**Figure S7.**
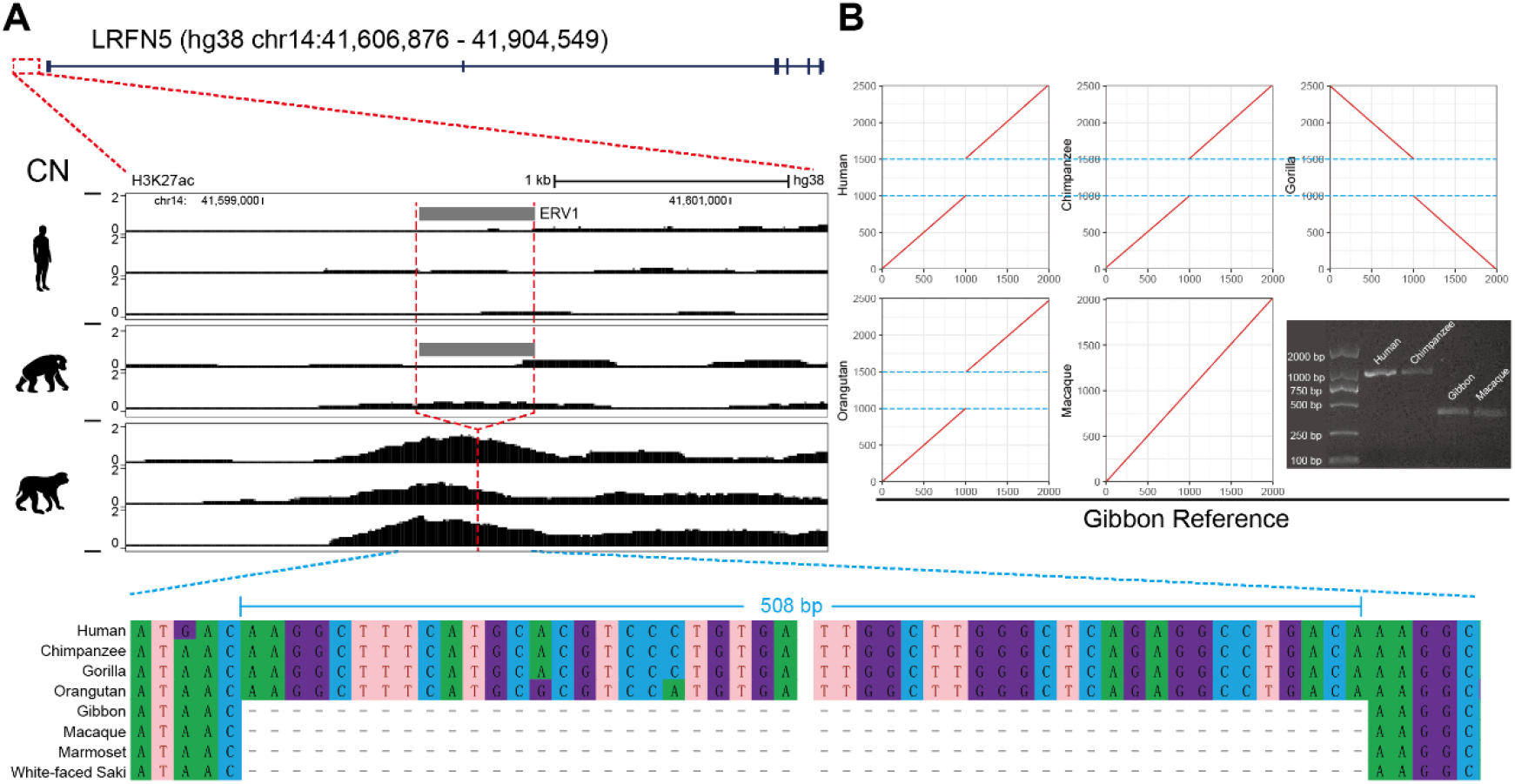
The data of the LRFN5 gene that contains a 508-bp insertion SV. (A) The H3K27ac signals among human, chimpanzee and macaque. The red dashed line indicates the position of SV. The DNA sequence alignment is present at the bottom. (B) The MUMMER plot and the result of PCR validation.

**Figure S8.**
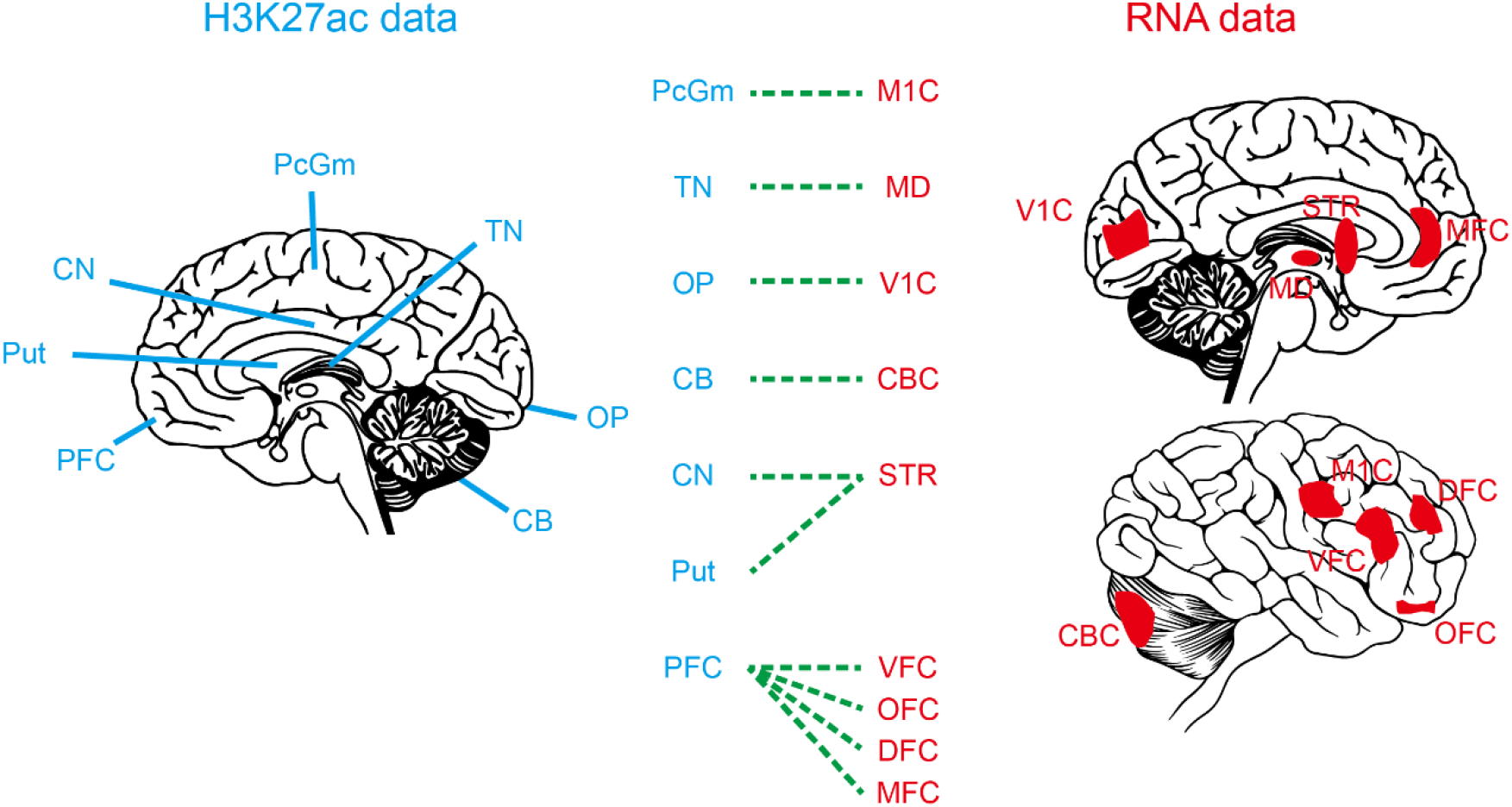
The matched brain regions between the H3K27ac data and the RNA-seq data.

**Figure S9.**
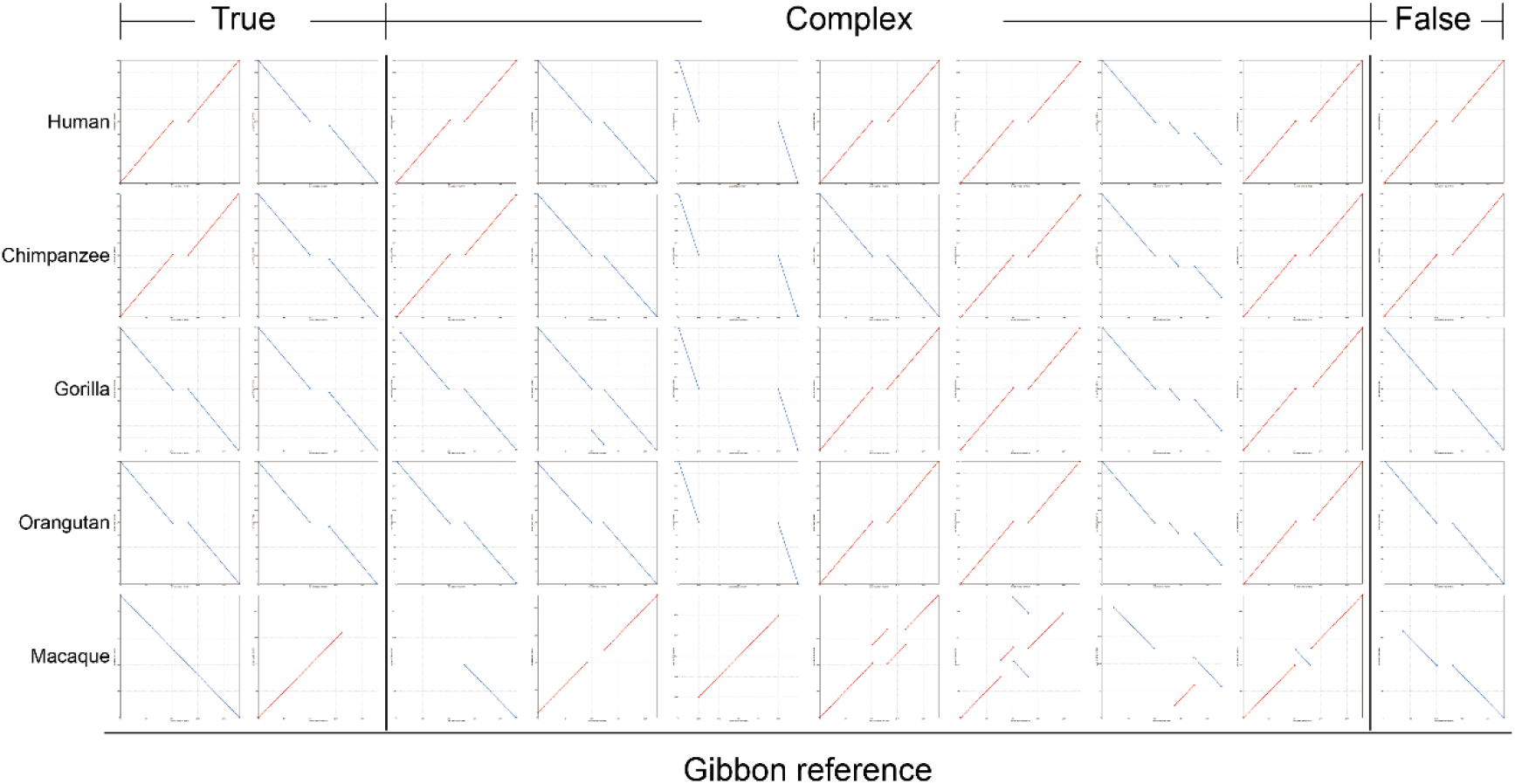
The MUMMER plots showing the patterns of the three defined DEL sets by manual checking. The three sets include the true DELs (defined as the DEL HC-GSSVs), the complex DELs and the false DELs. INSs are defined using the same strategy.

## Notes

### Competing Interest Statement

The authors have declared no competing interest.

## References

Abdalla, Ebtesam et al. 2022. “Cardiofacioneurodevelopmental Syndrome: Report of a Novel Patient and Expansion of the Phenotype.” American Journal of Medical Genetics. Part A 188(8): 2448–53.

Abel, Haley J. et al. 2020. “Mapping and Characterization of Structural Variation in 17,795 Human Genomes.” Nature 583(7814): 83–89.

Alba, David M. 2010. “Cognitive Inferences in Fossil Apes (Primates, Hominoidea): Does Encephalization Reflect Intelligence?” Journal of anthropological sciences = Rivista di antropologia: JASS 88: 11–48.

Austin-Tse, Christina et al. 2013. “Zebrafish Ciliopathy Screen Plus Human Mutational Analysis Identifies C21orf59 and CCDC65 Defects as Causing Primary Ciliary Dyskinesia.” American Journal of Human Genetics 93(4): 672–86.

Barton, Robert A., and Chris Venditti. 2017. “Rapid Evolution of the Cerebellum in Humans and Other Great Apes.” Current biology: CB 27(8): 1249–50.

Chaisson, Mark J. P. et al. 2015. “Resolving the Complexity of the Human Genome Using Single-Molecule Sequencing.” Nature 517(7536): 608–11.

Chen, X. et al. 2011. “GWA Study Data Mining and Independent Replication Identify Cardiomyopathy-Associated 5 (CMYA5) as a Risk Gene for Schizophrenia.” Molecular Psychiatry 16(11): 1117–29.

Chiang, Colby et al. 2017. “The Impact of Structural Variation on Human Gene Expression.” Nature Genetics 49(5): 692–99.

Chivers, D. J. 1998. “Measuring Food Intake in Wild Animals: Primates.” The Proceedings of the Nutrition Society 57(2): 321–32.

DeMari, Joseph et al. 2016. “CLTC as a Clinically Novel Gene Associated with Multiple Malformations and Developmental Delay.” American Journal of Medical Genetics. Part A 170A(4): 958–66.

Di Mattia, Thomas et al. 2018. “Identification of MOSPD2, a Novel Scaffold for Endoplasmic Reticulum Membrane Contact Sites.” EMBO reports 19(7): e45453.

Feng, Xiaowen, and Heng Li. 2021. “Higher Rates of Processed Pseudogene Acquisition in Humans and Three Great Apes Revealed by Long-Read Assemblies.” Molecular Biology and Evolution 38(7): 2958–66.

Gao, Yunhe et al. 2017. “Ring Finger Protein 43 Associates with Gastric Cancer Progression and Attenuates the Stemness of Gastric Cancer Stem-like Cells via the Wnt-β/Catenin Signaling Pathway.” Stem Cell Research & Therapy 8(1): 98.

Gordon, David et al. 2016. “Long-Read Sequence Assembly of the Gorilla Genome.” Science (New York, N.Y.) 352(6281): aae0344.

Goto-Ito, Sakurako et al. 2018. “Structural Basis of Trans-Synaptic Interactions between PTPδ and SALMs for Inducing Synapse Formation.” Nature Communications 9(1): 269.

Grant, Charles E., Timothy L. Bailey, and William Stafford Noble. 2011. “FIMO: Scanning for Occurrences of a given Motif.” Bioinformatics (Oxford, England) 27(7): 1017–18.

He, Yaoxi et al. 2019. “Long-Read Assembly of the Chinese Rhesus Macaque Genome and Identification of Ape-Specific Structural Variants.” Nature Communications 10(1): 4233.

Hill, Andrew, and Steven Ward. 1988. “Origin of the Hominidae: The Record of African Large Hominoid Evolution between 14 My and 4 My.” American Journal of Physical Anthropology 31(S9): 49–83.

Holloway, Ralph L., Chet C. Sherwood, Patrick R. Hof, and James K. Rilling. 2009. “Evolution of the Brain in Humans – Paleoneurology.” In Encyclopedia of Neuroscience, eds. Marc D. Binder, Nobutaka Hirokawa, and Uwe Windhorst. Berlin, Heidelberg: Springer Berlin Heidelberg, 1326–34. http://link.springer.com/10.1007/978-3-540-29678-2_3152 (January 4, 2023).

Hoya, Satoshi, Yuichiro Watanabe, Masako Shibuya, and Toshiyuki Someya. 2018. “Updated Meta-Analysis of CMYA5 Rs3828611 and Rs4704591 with Schizophrenia in Asian Populations.” Early Intervention in Psychiatry 12(5): 938–41.

Hu, Xuyun et al. 2017. “Novel Pathogenic ACAN Variants in Non-Syndromic Short Stature Patients.” Clinica Chimica Acta; International Journal of Clinical Chemistry 469: 126–29.

Infante, Jon et al. 2020. “POLR3A-Related Spastic Ataxia: New Mutations and a Look into the Phenotype.” Journal of Neurology 267(2): 324–30.

King, M. C., and A. C. Wilson. 1975. “Evolution at Two Levels in Humans and Chimpanzees.” Science (New York, N.Y.) 188(4184): 107–16.

Kronenberg, Zev N. et al. 2018. “High-Resolution Comparative Analysis of Great Ape Genomes.” Science (New York, N.Y.) 360(6393): eaar6343.

Le Duc, Diana et al. 2019. “Pathogenic WDFY3 Variants Cause Neurodevelopmental Disorders and Opposing Effects on Brain Size.” Brain: A Journal of Neurology 142(9): 2617–30.

Lee, Nam Soo et al. 2018. “Ring Finger Protein 126 (RNF126) Suppresses Ionizing Radiation-Induced P53-Binding Protein 1 (53BP1) Focus Formation.” The Journal of Biological Chemistry 293(2): 588–98.

Li, Heng. 2018. “Minimap2: Pairwise Alignment for Nucleotide Sequences.” Bioinformatics (Oxford, England) 34(18): 3094–3100.

Lin, Zhaohan et al. 2018. “Structural Basis of SALM5-Induced PTPδ Dimerization for Synaptic Differentiation.” Nature Communications 9(1): 268.

MacLeod, Carol E. et al. 2003. “Expansion of the Neocerebellum in Hominoidea.” Journal of Human Evolution 44(4): 401–29.

Morawski, M., G. Brückner, T. Arendt, and R. T. Matthews. 2012. “Aggrecan: Beyond Cartilage and into the Brain.” The International Journal of Biochemistry & Cell Biology 44(5): 690–93.

Morris, Mark R., Dewi Astuti, and Eamonn R. Maher. 2013. “Perlman Syndrome: Overgrowth, Wilms Tumor Predisposition and DIS3L2.” American Journal of Medical Genetics. Part C, Seminars in Medical Genetics 163C(2): 106–13.

Nabais Sá, Maria J. et al. 2020. “De Novo CLTC Variants Are Associated with a Variable Phenotype from Mild to Severe Intellectual Disability, Microcephaly, Hypoplasia of the Corpus Callosum, and Epilepsy.” Genetics in Medicine: Official Journal of the American College of Medical Genetics 22(4): 797–802.

Pan, Yaozhen et al. 2018. “POU5F1B Promotes Hepatocellular Carcinoma Proliferation by Activating AKT.” Biomedicine & Pharmacotherapy = Biomedecine & Pharmacotherapie 100: 374–80.

Panagopoulos, Ioannis, Emely Möller, Anna Collin, and Fredrik Mertens. 2008. “The POU5F1P1 Pseudogene Encodes a Putative Protein Similar to POU5F1 Isoform 1.” Oncology Reports 20(5): 1029–33.

Patel, Anand, Richard Schwab, Yu-Tsueng Liu, and Vineet Bafna. 2014. “Amplification and Thrifty Single-Molecule Sequencing of Recurrent Somatic Structural Variations.” Genome Research 24(2): 318–28.

Pozzi, Luca et al. 2014. “Primate Phylogenetic Relationships and Divergence Dates Inferred from Complete Mitochondrial Genomes.” Molecular Phylogenetics and Evolution 75: 165–83.

Rafnar, Thorunn et al. 2014. “Genome-Wide Association Study Yields Variants at 20p12.2 That Associate with Urinary Bladder Cancer.” Human Molecular Genetics 23(20): 5545–57.

Roth, Gerhard, and Ursula Dicke. 2012. “Evolution of the Brain and Intelligence in Primates.” Progress in Brain Research 195: 413–30.

Russell, Jonathan F. et al. 2012. “Familial Cortical Myoclonus with a Mutation in NOL3.” Annals of Neurology 72(2): 175–83.

Simó-Riudalbas, Laia et al. 2022. “Transposon-Activated POU5F1B Promotes Colorectal Cancer Growth and Metastasis.” Nature Communications 13(1): 4913.

Smith, R. J., and W. L. Jungers. 1997. “Body Mass in Comparative Primatology.” Journal of Human Evolution 32(6): 523–59.

Sousa, André M. M. et al. 2017. “Molecular and Cellular Reorganization of Neural Circuits in the Human Lineage.” Science 358(6366): 1027–32.

Spielmann, Malte, Darío G. Lupiáñez, and Stefan Mundlos. 2018. “Structural Variation in the 3D Genome.” Nature Reviews. Genetics 19(7): 453–67.

Stankiewicz, Pawel, and James R. Lupski. 2010. “Structural Variation in the Human Genome and Its Role in Disease.” Annual Review of Medicine 61: 437–55.

Suntsova, Maria V., and Anton A. Buzdin. 2020. “Differences between Human and Chimpanzee Genomes and Their Implications in Gene Expression, Protein Functions and Biochemical Properties of the Two Species.” BMC genomics 21(Suppl 7): 535.

Suo, Guangli et al. 2005. “Oct4 Pseudogenes Are Transcribed in Cancers.” Biochemical and Biophysical Research Communications 337(4): 1047–51.

Vermunt, Marit W. et al. 2016. “Epigenomic Annotation of Gene Regulatory Alterations during Evolution of the Primate Brain.” Nature Neuroscience 19(3): 494–503.

Warren, Wesley C. et al. 2020. “Sequence Diversity Analyses of an Improved Rhesus Macaque Genome Enhance Its Biomedical Utility.” Science (New York, N.Y.) 370(6523): eabc6617.

Watanabe, H., Y. Yamada, and K. Kimata. 1998. “Roles of Aggrecan, a Large Chondroitin Sulfate Proteoglycan, in Cartilage Structure and Function.” Journal of Biochemistry 124(4): 687–93.

Wei, Ming et al. 2021. “Identification of Novel ACAN Mutations in Two Chinese Families and Genotype-Phenotype Correlation in Patients with 74 Pathogenic ACAN Variations.” Molecular Genetics & Genomic Medicine 9(11): e1823.

Weischenfeldt, Joachim, Orsolya Symmons, François Spitz, and Jan O. Korbel. 2013. “Phenotypic Impact of Genomic Structural Variation: Insights from and for Human Disease.” Nature Reviews. Genetics 14(2): 125–38.

Wheatley, Bruce P. 1987. “The Evolution of Large Body Size in Orangutans: A Model for Hominoid Divergence.” American Journal of Primatology 13(3): 313–24.

Wu, Tianzhi et al. 2021. “ClusterProfiler 4.0: A Universal Enrichment Tool for Interpreting Omics Data.” Innovation (Cambridge (Mass.)) 2(3): 100141.

Zanke, Brent W. et al. 2007. “Genome-Wide Association Scan Identifies a Colorectal Cancer Susceptibility Locus on Chromosome 8q24.” Nature Genetics 39(8): 989–94.

Zihlman, A. L., and R. K. McFarland. 2000. “Body Mass in Lowland Gorillas: A Quantitative Analysis.” American Journal of Physical Anthropology 113(1): 61–78.

Zihlman, Adrienne L., and Debra R. Bolter. 2015. “Body Composition in Pan Paniscus Compared with Homo Sapiens Has Implications for Changes during Human Evolution.” Proceedings of the National Academy of Sciences of the United States of America 112(24): 7466–71.

